# Whole-genome sequence analysis of mutations in rice plants regenerated from zygotes, mature embryos, and immature embryos

**DOI:** 10.1101/2021.01.20.427397

**Authors:** Masako Ichikawa, Norio Kato, Erika Toda, Masakazu Kashihara, Yuji Ishida, Yukoh Hiei, Sachiko N. Isobe, Kenta Shirasawa, Hideki Hirakawa, Takashi Okamoto, Toshihiko Komari

## Abstract

Somaclonal variation was studied by whole-genome sequencing in rice plants (*Oryza sativa* L., ‘Nipponbare’) regenerated from the zygotes, mature embryos, and immature embryos of a single mother plant. The mother plant and its seed-propagated progeny were also sequenced. A total of 338 variants of the mother plant sequence were detected in the progeny, and mean values ranged from 9.0 of the seed-propagated plants to 37.4 of regenerants from mature embryos. The ratio of single nucleotide variants among the variants was 74.3%, and the natural mutation rate calculated using the variants in the seed-propagated plants was 1.2 × 10^−8^. The percentage and the mutation rate were consistent with the values reported previously. Plants regenerated from mature embryos had significantly more variants than different progeny types. Therefore, using zygotes and immature embryos can reduce somaclonal variation during the genetic manipulation of rice.

---

Cell culture enables foreign gene transfer and gene editing in plant science. Genetic modifications are first introduced into a single cell, and then plants are regenerated from culturing that modified cell. However, previous research has shown that heritable changes, known as somaclonal variation (Nishi et al. 1968, Heinz and Mee 1969, Larkin and Scowcroft 1981), frequently occur in regenerants of dedifferentiated cells. The characteristics and induction mechanisms of somaclonal variation are poorly understood. Some variation is attributed to differences in DNA methylation levels between cells that propagate *in vivo* and *in vitro*. For example, mutations caused by transposable elements are normally suppressed *in vivo* by DNA methylation but are induced in cultured cells (Hirochika et al. 1996). Prolonged culturing leads to higher mutation rates (Qin et al. 2018), and so it is typically avoided in gene transfer, editing, and other cell engineering practices. However, when somaclonal mutations are desired (e.g., to introduce more variation in breeding populations), prolonged culturing can be useful (Larkin and Scowcroft 1981).

Studying genetic variation in regenerants from cell cultures, transgenic plants, and edited plants by whole-genome sequencing has become feasible because of recent DNA sequencing technology advancements. Rice (*Oryza sativa* L.) is one of the most frequently studied plant species. Somaclonal variation in rice plants, including transgenic and edited plants, has been analyzed by whole-genome sequencing (Miyao et al. 2012, Kawakatsu et al. 2013, Zhang et al. 2014, Wei et al. 2016, Qing et al. 2018, Tang et al. 2018, and Park et al. 2019). Such studies detected numerous mutations that were induced by cell culture. Varietal differences in somaclonal variation in rice have also been observed with amplified fragment length polymorphism (AFLP) analysis (Wang et al. 2013).

One limitation of the studies cited above is that the sequenced plants were regenerated solely from mature embryos. Other cell and tissue sources should be investigated to gain a better understanding of somaclonal variation. For example, immature embryos may be more efficient for genetic transformation in cereal plants (Hiei et al. 2014). More efficient gene editing has been reported in rice zygotes (Toda et al. 2019). Another limitation of the studies cited above is that cells were cultured for more than two months in most cases, although a shorter culture period is typically recommended. Toki (1997) described a rice transformation protocol that requires only one month to grow calli from mature seeds. Therefore, in this study, rice plants were regenerated from zygotes, mature embryos, and immature embryos using the minimal times required to achieve callus induction and growth. The mutation rates of regenerants were compared by whole-genome resequencing.

The process for obtaining regenerated plant samples is shown in Figure 1. A mother plant was cultivated in a greenhouse from a single seed of the rice cultivar ‘Nipponbare’ obtained from a commercial distributor. The following samples were collected from the mother plant and used immediately: young spikelets (before flowering), immature seeds (12 days after pollination), and mature seeds.

**Figure 1.**
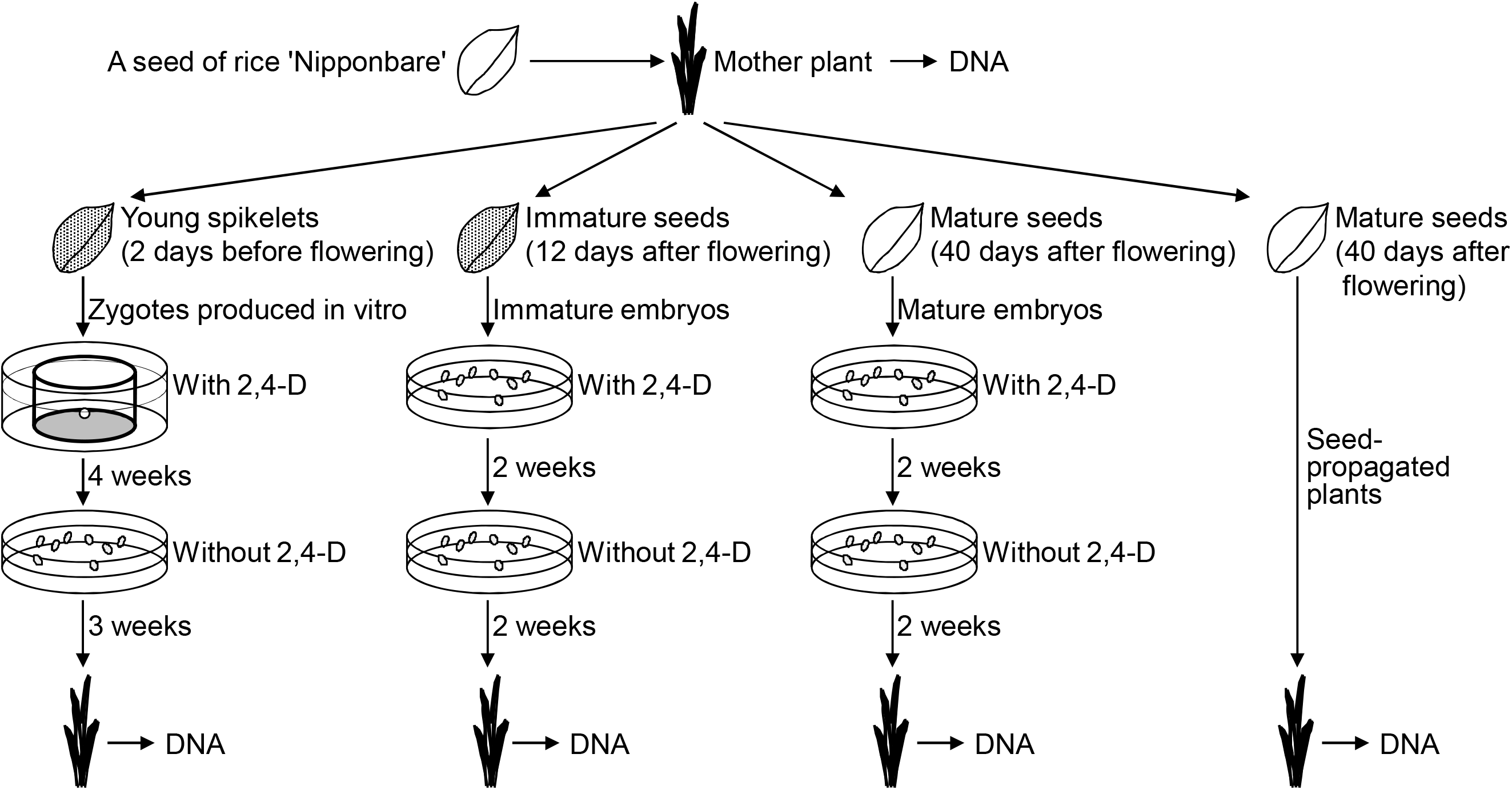
Sampling design for whole-genome sequencing. DNA was isolated from 5 plants derived from 5 different spikelets/seeds in each progeny group.

Egg and sperm cells were isolated from young spikelets and electrically fused to produce zygotes, as described by Uchiumi et al. (2006, 2007). The zygotes were cultured and regenerated into plantlets (Uchiumi et al. 2007). Husking and sterilizing mature seeds, preparing immature embryos from immature seeds, and making culture media to induce calli and regenerate plants from mature and immature embryos were performed as previously described (Hiei et al. 2008). Callus induction and cell culture from zygotes, immature embryos, and mature embryos using media that contains 2, 4-dichlorophenoxy-acetic acid (2, 4-D) were conducted for 4, 2, and 2 weeks, respectively. Periods for the plant regeneration cultures without 2, 4-D from the cells from zygotes, immature embryos, and mature embryos were 3, 2, and 2 weeks, respectively. Regenerants were grown in a greenhouse with seed-propagated plants.

The plants shown in Figure 1 were analyzed by whole-genome resequencing. No abnormalities were visually observed in greenhouse-grown plants. Approximately 1 g of young leaf segments were sampled from the mother plant and four progeny groups: (i) five regenerants from five independent zygotes, (ii) five regenerants from five independent immature embryos, (iii) five regenerants from five independent mature embryos, and (iv) five seed-propagated plants. The leaves were frozen and stored at −80 °C before DNA preparation.

Total DNA was extracted from the young leaves of 21 plants using a DNeasy Plant Mini Kit (Qiagen, Hilden, Germany). DNA quality was analyzed using NanoDrop™ and Qubit™ (Thermo Scientific, Waltham, Massachusetts, US). Illumina paired-end (PE) libraries were constructed for each plant with a TruSeq DNA PCR-Free Sample Prep Kit (Illumina, San Diego, CA). Illumina HiSeq X with a read length of 151 nucleotides (nt) was used to obtain genome sequences. Sequenced bases with quality scores less than 10 were filtered by PRINSEQ 0.20.4 (Schmieder and Edwards, 2011). Adaptor sequences in the reads were trimmed using FASTX-Clipper from the FASTX-Toolkit 0.0.13 (http://hannonlab.cshl.edu/fastx_toolkit).

A total of 3,066,032,798 reads were obtained from the 21 Illumina PE libraries. The average number of obtained reads per line was 146,001,562 (range: 59,349,518 to 188,539,542). The total lengths of the obtained reads per plant are shown in Figure 2. The Illumina PE reads were mapped onto the ‘Nipponbare’ reference genome (Os-Nipponbare-Reference-IRGSP-1.0, Kawahara et al. 2013) to detect base variances (SNVs and indels) by Bowtie2 (Langmead and Salzberg, 2012). Variant calls were performed by bcftools 1.9 in SAMtools (Li et al. 2009). The parameters used for variant calls were bcftools-1.9/bin/bcftools mpileup -d 10000000 -Ou -b bam-list.txt-f IRGSP-1.0_genome.fasta -a DP,AD,INFO/AD | bcftools-1.9/bin/bcftools call -cv -Ov -f GQ -o output.vcf.

**Figure 2.**
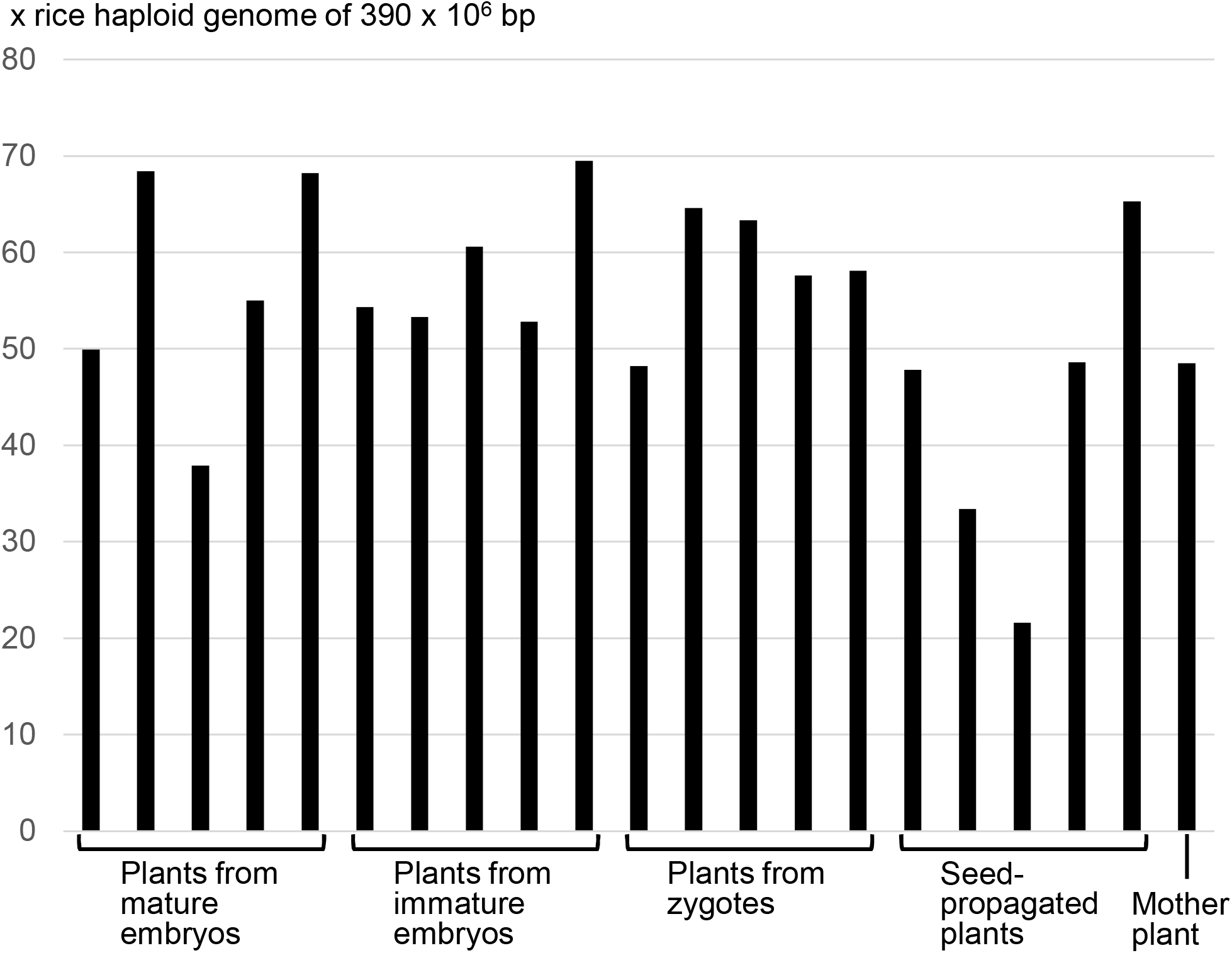
Total length of sequence reads per plant.

Variants of the mother plant were filtered to remove false positives using the following criteria: minimum depth = 10; maximum depth = 65; minimum quality = 999; variants at positions where bases were not called in all plants were excluded; variants that were detected in two or more plants at the same position were excluded. The number of variants of the mother plant in each progeny group was statistically compared, assuming Poisson distributions. Namely, when two groups were compared, the variant numbers in the two groups were hypothesized to be samples from the same Poisson distribution, and then the probability for such a combination of Poisson samples was calculated to test the hypothesis. Mutation rates per rice diploid genome were calculated by dividing the number of variants by 2 × 390 × 10^6^, as shown in Tang et al. (2018).

A total of 338 variants of the mother plant were detected in the progeny groups, and the number of the variants in each progeny group is shown in Figure 3. Mean values ranged from 9.0 of the seed-propagated plants to 37.4 of regenerants from mature embryos. The ratio of single nucleotide variants among the variants of the mother plant was 74.3%. This percentage is close to those reported by Zhang et al. (2014) and Tang et al. (2018). The natural mutation rate calculated using the variants in the seed-propagated plants was 1.2 × 10^−8^, which is consistent with the natural mutation rates of 5.4 × 10^−8^ for rice (Tang et al. 2018), 0.7 × 10^−8^ for Arabidopsis (Ossowski et al. 2010), and 2.2–3.9 × 10^−8^ for maize (Yang et al. 2017).

**Figure 3.**
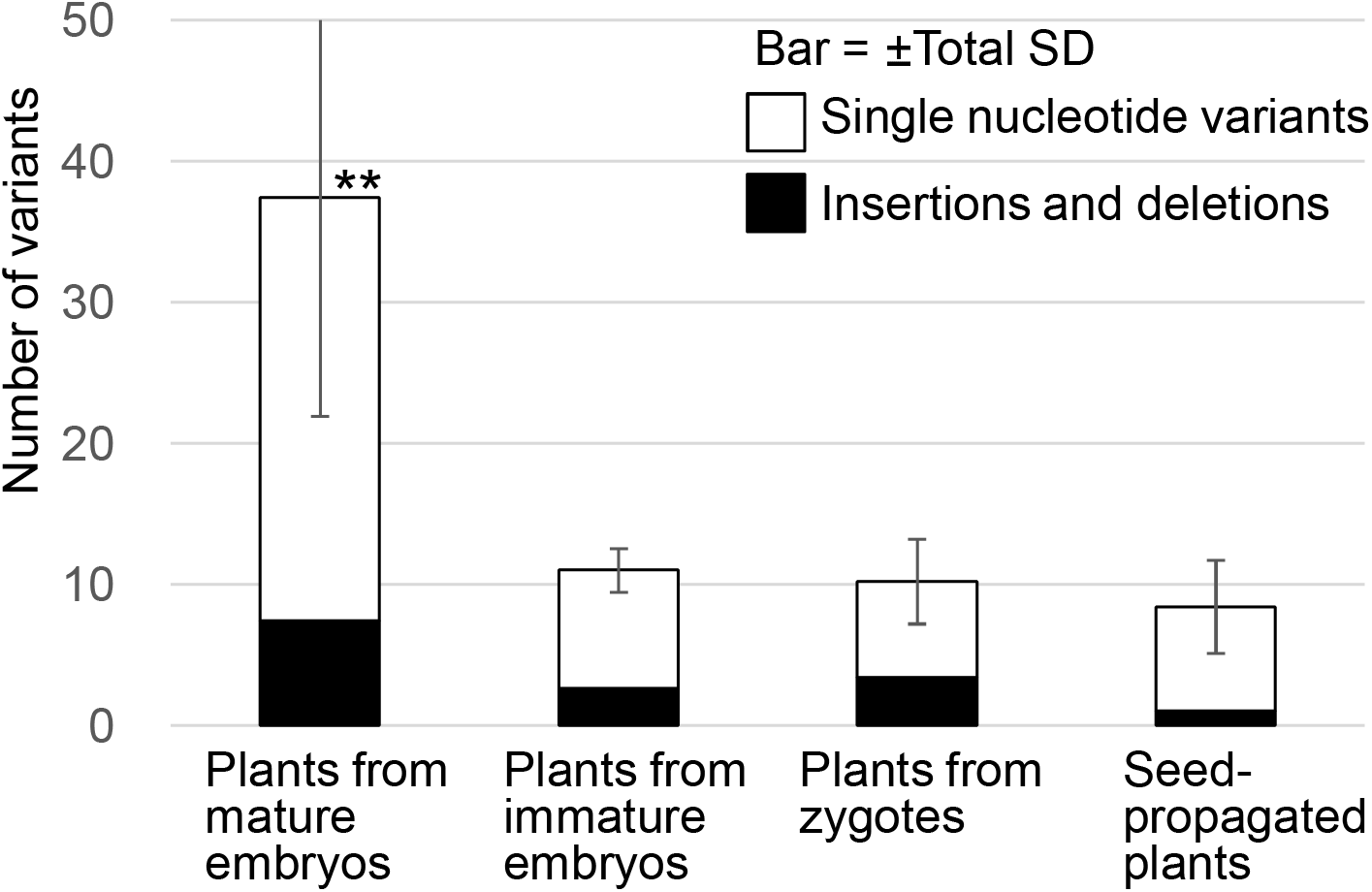
Number of variants of the mother plant. ** Significantly different from the other three progeny groups (P<0.01).

More variants were detected in regenerants from mature embryos than in regenerants from immature embryos or zygotes. The numbers of variants detected in regenerants from immature embryos and zygotes were similar to those detected in seed-propagated plants. These results suggest that the starting tissue affects mutation rates. The regenerants from zygotes were cultured longer than regenerants from other tissues because callus formation from single cells takes longer. Despite its extended duration, the process of culturing cells from zygotes was not more mutagenic than culturing cells from other tissues. The difference in the number of variants between regenerants cultured from mature and immature embryos was unexpected because calli induced from these tissues are similar in appearance. A callus is a mass of dedifferentiated cells and is generally believed to have lost the characteristics of the original tissue. More detailed comparisons, such as analyzing patterns of transcription and DNA methylation in calli from different tissues, may provide useful insight into the nature of somaclonal variation.

Mature seeds are commonly used and are a convenient starting tissue for cell culture because they can be obtained in large quantities and stored at room temperature for several months before culturing. In contrast, cell culture protocols from zygotes and immature embryos require a constant supply of fresh materials from healthy plants grown in well-conditioned greenhouses. However, the results of this study suggest that the use of immature embryos and zygotes should be added to the general guidance of shorter cell culture periods to reduce somaclonal variation during the genetic manipulation of rice.

## Abbreviations

SNV: single nucleotide variant
indels: insertions and deletions

## Acknowledgments

The authors thank Naoki Takemori for his support and helpful discussions and Chitose Matsui and Natsuki Hayashi for their technical assistance.

